# Glial-derived mitochondrial signals impact neuronal proteostasis and aging

**DOI:** 10.1101/2023.07.20.549924

**Authors:** Raz Bar-Ziv, Naibedya Dutta, Adam Hruby, Edward Sukarto, Maxim Averbukh, Athena Alcala, Hope R. Henderson, Jenni Durieux, Sarah U. Tronnes, Qazi Ahmad, Theodore Bolas, Joel Perez, Julian G. Dishart, Matthew Vega, Gilberto Garcia, Ryo Higuchi-Sanabria, Andrew Dillin

## Abstract

The nervous system plays a critical role in maintaining whole-organism homeostasis; neurons experiencing mitochondrial stress can coordinate the induction of protective cellular pathways, such as the mitochondrial unfolded protein response (UPR^MT^), between tissues. However, these studies largely ignored non-neuronal cells of the nervous system. Here, we found that UPR^MT^ activation in four, astrocyte-like glial cells in the nematode, *C. elegans*, can promote protein homeostasis by alleviating protein aggregation in neurons. Surprisingly, we find that glial cells utilize small clear vesicles (SCVs) to signal to neurons, which then relay the signal to the periphery using dense-core vesicles (DCVs). This work underlines the importance of glia in establishing and regulating protein homeostasis within the nervous system, which can then impact neuron-mediated effects in organismal homeostasis and longevity.

**One-Sentence Summary:** Glial cells sense mitochondrial stress and signal a beneficial stress signal to promote neuronal health and longevity.

## Main Text

Aging is a complex and gradual process accompanied by a progressive decline in physiological integrity and function. Underlying this physiological decline is the loss of essential cellular functions, concomitant with the accumulation of cellular and molecular damage, such as the increase in DNA mutations and accumulation of misfolded proteins, causing a decline in organellar and cellular function (*1*, *2*). One of the key organ systems in aging is the nervous system. The nervous system is not only impacted by aging, as exemplified by neurodegenerative diseases, but can also modulate the aging process of the entire organism (*3*). The link between the nervous system and aging is governed by its role as a central regulatory hub, mediating the maintenance of homeostasis in response to fluctuations in the external and internal environments, intracellularly and intercellularly, within and between different tissues. One level of homeostasis that neurons are tasked with maintaining and coordinating across tissues is the homeostatic state of protein folding (proteostasis, protein homeostasis) (*4*). Proteostasis is one of the key features declining with age, due to the reduced ability to counteract misfolding and aggregation of proteins, a hallmark of many neurodegenerative diseases, such as Huntington’s disease (*5*). Neurons are able to regulate protein homeostasis across tissues for several pathways that are dedicated to protecting different cellular compartments (*6*, *7*), such as the mitochondrion (*8*), with activation of the pathway prolonging lifespan, by coordinating mitochondrial protein homeostasis across tissues. This protective pathway, the unfolded protein response of mitochondria (UPR^MT^) (*9*), is triggered upon insults to the organelle and causes the activation of a coordinated transcriptional response, which includes the upregulation of mitochondrially-targeted chaperones and proteases, modulating cellular processes to alleviate the burden of stress on mitochondria. Interestingly, neurons secrete molecules that allow coordination of the UPR^MT^ across tissues, utilizing dense-core vesicles (DCVs), serotonin, and the WNT-like ligand EGL-20 to trigger a response in peripheral tissues, such as the intestine (*10*, *11*).

Historically, much scientific work interrogating the homeostatic roles of the nervous system focused on neurons. While it is clear that glia, the other main cell type of the nervous system, can serve many roles in neuronal development and function, these roles are normally associated with support roles, including regulating cell number, neuronal migration, axon specification and growth, synapse formation and pruning, ion homeostasis, synaptic plasticity, and providing metabolic support for neurons (*12*, *13*). However, in recent years, it has become increasingly clear that glial health can impact aging and progression of neurodegenerative diseases, like Alzheimer’s Disease (AD) (*14*). For example, expression of ApoE4, one of the strongest risk factors for AD, specifically in astrocytes resulted in increased neuronal tau aggregation (*15*). Moreover, hyperactivation of the unfolded protein response of the endoplasmic reticulum (UPR^ER^), which drives ER stress resilience, solely in astrocyte-like glial cells resulted in a significant lifespan extension in *C. elegans* (*16*). While these studies show the importance of glial function in organismal health, what they lacked is an active function of glia in promoting these beneficial effects. To uncover an active role for glia in stress signaling and longevity, we aimed to determine whether glial cells can sense mitochondrial stress and initiate an organism-wide response to promote mitochondrial stress resilience and longevity. We used multiple genetic methods to activate UPR^MT^ in non-neuronal cells, including cell-type specific application of mitochondrial stress and direct activation of the UPR^MT^ in the absence of stress. Interestingly, we found that regardless of method, activation of UPR^MT^ in a small subset of glial cells, the cephalic sheath (CEPsh) glia, provided robust organismal benefits, including prolonged lifespan and increased resistance to oxidative stress. Perhaps most unique in this model is that UPR^MT^ activation in CEPsh glia promotes neuronal health by alleviating protein aggregation in neurons of a Huntington’s disease model. In fact, CEPsh glia directly communicate with neurons through the release of small clear vesicles (SCVs) and relay the coordination to the periphery via downstream neuronal mechanisms. This glia to neuron signal results in induction of the UPR^MT^ in distal tissues, through a cell non-autonomous mechanism, which is dependent on the canonical UPR^MT^ pathway, yet surprisingly distinct from paradigms where UPR^MT^ is directly activated in neurons. Collectively, these results reveal a novel function for CEPsh glia in sensing mitochondrial stress, which initiates a signal to promote protein homeostasis in neurons and ultimately prolongs longevity. Therefore, glial cells serve as one of the upstream mediators of mitochondrial stress and its coordination across the entire organism.

## RESULTS

### Activation of the UPR^MT^ in glia elicits beneficial effects for the entire organism

To activate the UPR^MT^ in glial cells, we leveraged the ability of JMJD-1.2 (JumonjiC (JmjC)-domain-containing protein), a histone demethylase, to induce a robust UPR^MT^ in the absence of stress (*17*). Specifically, we overexpressed *jmjd-1.2* under several established glial promoters (*18*), targeting most glial cells, or glial sub-types using specific promoters. The activation of UPR^MT^ in most glial cells resulted in a mild lifespan extension (**Fig. 1A and fig. S1A**). Interestingly, we found that overexpressing *jmjd-1.2* in either amphid sheath and phasmid sheath glia (*fig-1p*), or the four CEPsh glia (*hlh-17p*) alone was not only sufficient to prolong lifespan, but had a more profound effect than overexpressing *jmjd-1.2* in most glial cells simultaneously (**Fig. 1, B and C**). On the cellular level, the activation of the UPR^MT^ in all different glial subtypes was also able to trigger the activation of the UPR^MT^ in distal, intestinal cells, as can be observed by the activation of a fluorescent reporter under the regulation of a mitochondrial chaperone promoter (*hsp-6p::GFP*) (**Fig. 1, D and E**). Similar to lifespan extensions, overexpression of *jmjd-1.2* in CEPsh glia had the most profound effect on distal UPR^MT^ activation. We confirm that this is a beneficial, functional activation of UPR^MT^ as these animals also show increased resistance to paraquat (**Fig. 1F and fig. S1G**), a drug that increases superoxide levels mainly in the mitochondria (*19*).

**Figure 1:**
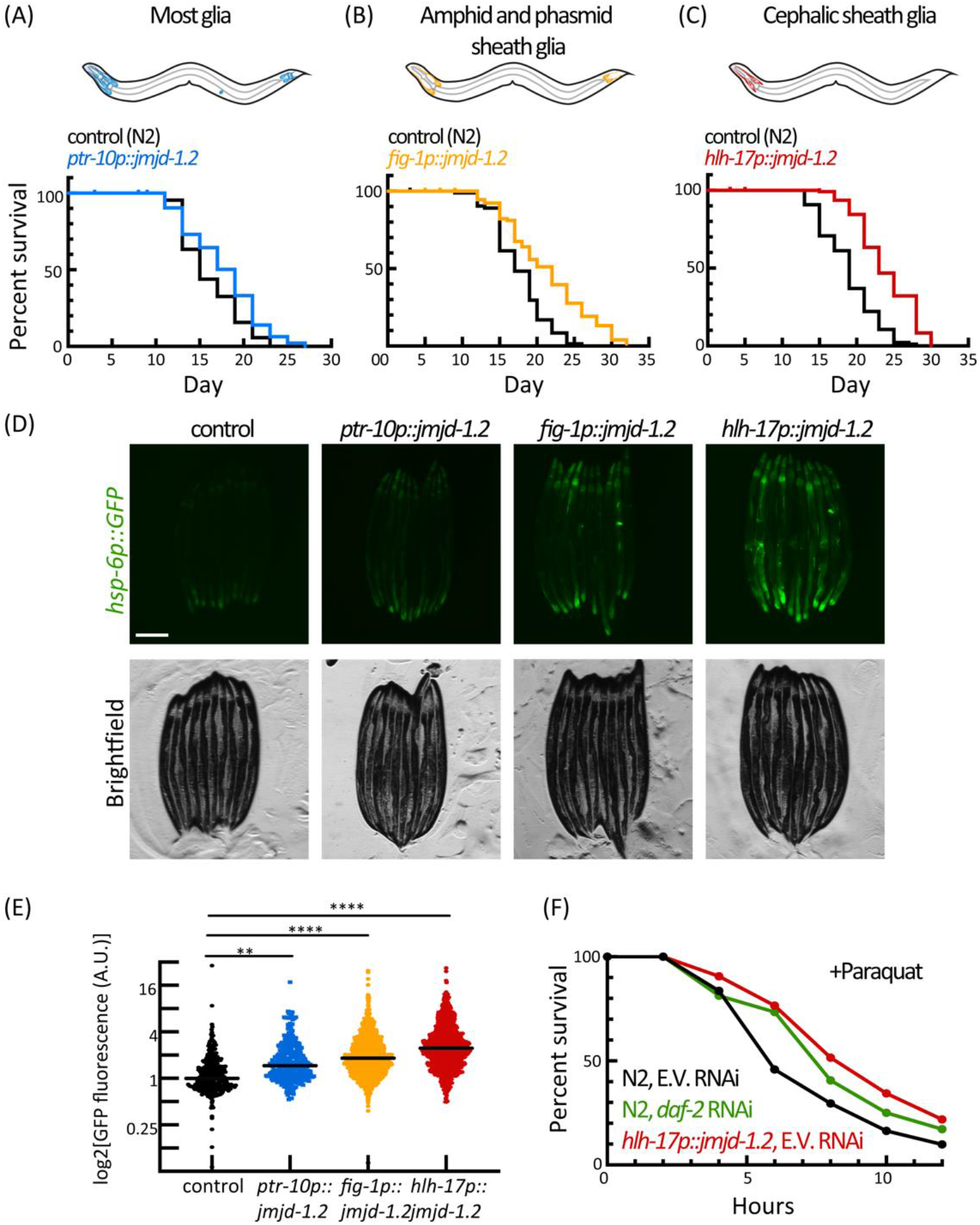
Glial activation of *jmjd-1.2* prolongs lifespan, improves stress resistance, and induces the cell non-autonomous UPR^MT^. (A) Survival of animals expressing *jmjd-1.2* in most glia (blue), compared to a control N2 population (black). P= 0.0158. (B) Survival of animals expressing *jmjd-1.2* in amphid and phasmid sheath glia (orange), compared to a control N2 population. Log-rank test, P < 0.0001 (C) Survival of animals expressing *jmjd-1.2* in the cephalic sheath (CEPsh) glia (red), compared to a control N2 population. P < 0.0001 (D) Representative fluorescent micrographs of UPR^MT^ reporter worms (*hsp-6p::GFP*) expressing *jmjd-1.2* under the indicated promoters. Scale bar, 250 μm. (E) Fold change in GFP fluorescence per worm using a large-particle biosorter (methods), normalized to a control (*hsp-6p::GFP*) population (n>300 per group). One-way analysis of variance (ANOVA) Tukey’s multiple comparisons test, **P < 0.01, ****P < 0.0001. See also Figure S1G. (F) Survival of animals expressing glial *jmjd-1.2* (red) under paraquat stress, as compared to a control population (black). Worms fed with *daf-2* RNAi were used as a positive control (green) (n=60 per group), see also Figure S1H.

The most profound peripheral UPR^MT^ activation and lifespan extension was observed when overexpressing *jmjd-1.2* in CEPsh glia, the four astrocyte-like cells that associate with sensory organs and extend processes that wrap around the nerve ring, the major neuropil of *C. elegans* (*12*). Therefore, we focused our efforts on this specific glial subtype (heretofore referred to as glial *jmjd-1.2*). To determine whether CEPsh glia can elicit non-autonomous signals in direct response to stress rather than solely via ectopic overexpression of UPR^MT^ activators, we induced mitochondrial stress specifically in CEPsh glia in two different ways: 1) expression of a mitochondrially-targeted KillerRed to induce localized oxidative stress in mitochondria, and 2) expression of the mitochondrial-binding polyglutamine tract Q40 (*10*). The introduction of mitochondrial stress in CEPsh glia using both methods resulted in a similar activation of UPR^MT^ in peripheral tissue as *jmjd-1.2* overexpression (**fig. S1, B to D**). Furthermore, we verified our results using an alternative CEPsh glia-specific promoter recently identified using a single-cell RNA-seq dataset (**fig. S1E**) (*20*). Lastly, the distal activation by CEPsh glia was dependent on intact glial cells, as ablating the cells using the mutant *vab-3* (*21*) abrogated the effect (**fig. S1F**). Taken together, these data indicate that not only can CEPsh glia communicate mitochondrial stress in a non-autonomous manner, but they can directly sense mitochondrial stress. Importantly, these studies also highlight the utility of the *jmjd-1.2* overexpression line as a viable mimetic for non-autonomous communication downstream of mitochondrial stress in glial cells without the caveats of the potentially detrimental effects of mitochondrial dysfunction in neural cells (*10*).

We sought to further decipher the underlying mechanisms of the cell non-autonomous activation triggered by CEPsh glia. First, we tested whether the UPR^MT^ transcriptional program mediated by DVE-1/UBL-5 is also induced upon glial UPR^MT^ induction (*22*, *23*). DVE-1 is a transcription factor, acting with UBL-5, and mediating UPR^MT^ to alleviate stress in the mitochondria (*22*). We found that inducing the UPR^MT^ in CEPsh glia caused an increase in the DVE-1::GFP signal both in the head and the intestinal region of the worm (**Fig. 2, A and B**). Using the fluorescent reporter *hsp-6p::GFP*, we also found that the activation of the distal UPR^MT^ in response to glial signals depends on three different transcriptional programs of the UPR^MT^: ATFS-1, UBL-5/DVE-1, and LIN-65/MET-2 (*24*) (**Fig. 2, C to E and fig. S2**). Intact UPR^MT^ was also required for the longevity phenotype, as knocking down the transcription factor ATFS-1, which suppressed UPR^MT^ activation, also abrogated the lifespan extension of the glial *jmjd-1.2* animals (**Fig. 2F**). Collectively, these results indicate that the canonical components of the UPR^MT^ pathway are required for the peripheral response of cell non-autonomous communication from CEPsh glia.

**Figure 2:**
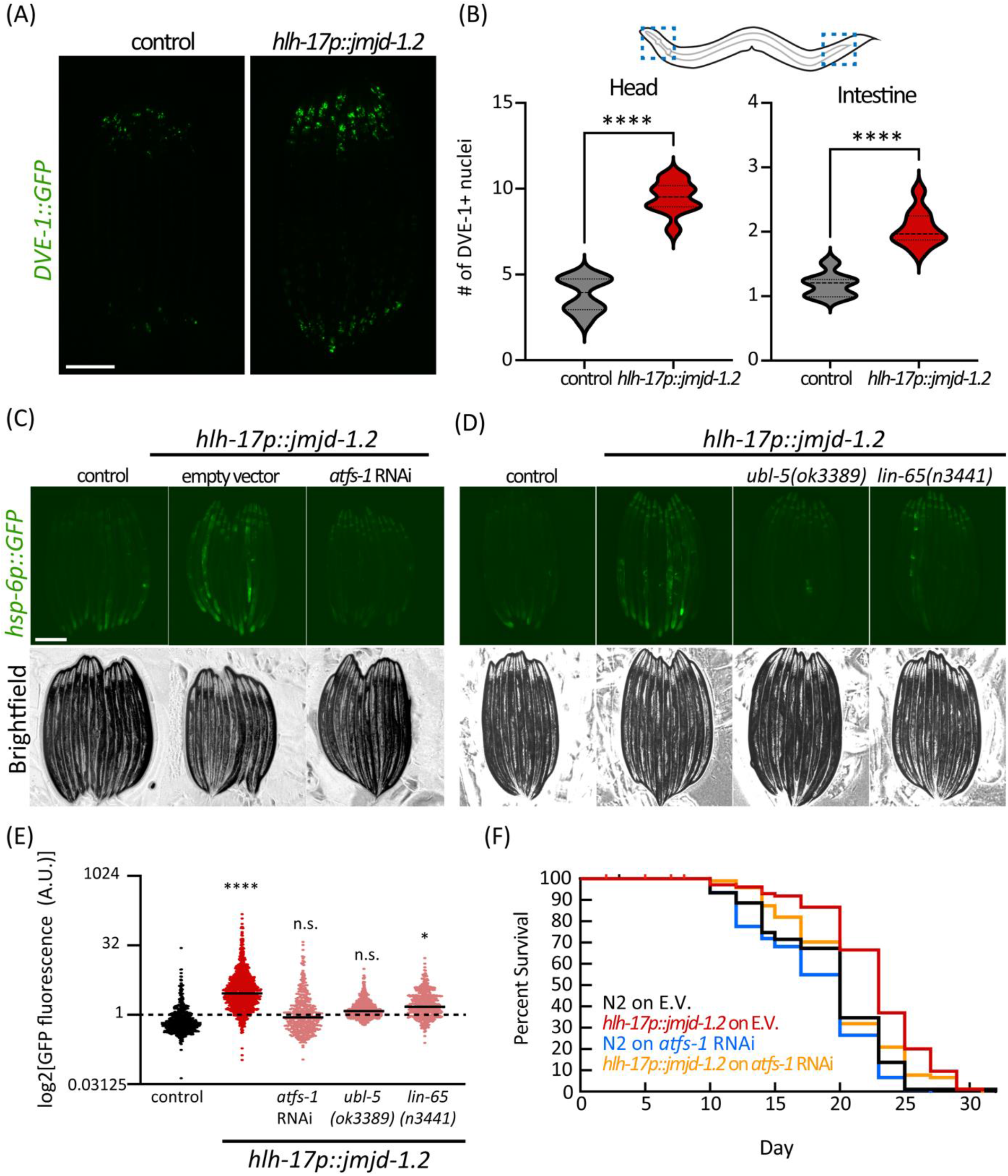
Non-autonomous activation of UPR^MT^ in the periphery depends on cell-autonomous regulators of the pathway UPR^MT^. (A) Representative fluorescent micrograph of UPR^MT^ translational reporter worms (*DVE-1::GFP*) expressing *jmjd-1.2* under the CEPsh glia (*hlh-17)* promoter, and the number of DVE-1+ nuclei quantified for the anterior and posterior regions using Fiji (n>30). Scale bar, 250 μm. (B), significance was assessed relative to control by unpaired Student’s t test, ****P<0.0001. (C and D) Representative fluorescent micrograph of UPR^MT^ reporter worms (*hsp-6p::GFP*) expressing *jmjd-1.2* either knocked-down (C) or knocked-out (D) for key regulators in UPR^MT^ activation, and quantified in (E), as in Figure 1E, (n>450), One-way analysis of variance (ANOVA) Tukey’s multiple comparisons test. n.s. not significant, *P<0.05, ****P < 0.0001. See also Figure S2 (F) Survival of animals expressing *jmjd-1.2* in CEPsh glia grown on either control (E.V.) or *atfs-1* RNAi.

### Activation of UPR^MT^ in glia rewires multiple cellular processes

To examine more global changes that occur upon activation of the UPR^MT^ in CEPsh glia, we profiled gene expression changes using whole-animal bulk RNA-seq. We observed many changes in gene expression in glial *jmjd-1.2* animals with 547 significantly upregulated genes and 413 downregulated genes compared to a wild-type control (**Fig. 3A**). As expected, gene expression changes show large similarities to other UPR^MT^ paradigms including electron transport chain (ETC) inhibition via *cox-5B* RNAi (*25*) or whole-animal overexpression of *jmjd-1.2* (*17*) (**Fig. 3B**). Indeed, animals with glial *jmjd-1.2* overexpression showed an overall increase in UPR^MT^ genes (**Fig. 3, C and D**), while other stress pathways did not show a significant difference. Moreover, we observed a decrease in the expression of genes involved in translation and ribosome biology (**Fig. 3C**), a characteristic of stress responses meant to alleviate the protein burden on the organelle (*26*). We also observed an overrepresentation of genes involved in the response to oxidative stress, with ∼40% of the transcriptional response overlapping with the response to paraquat (**Fig. 3E**), consistent with our data demonstrating that animals with *jmjd-1.2* induction in CEPsh glia exhibit increased resilience to oxidative stress.

**Figure 3:**
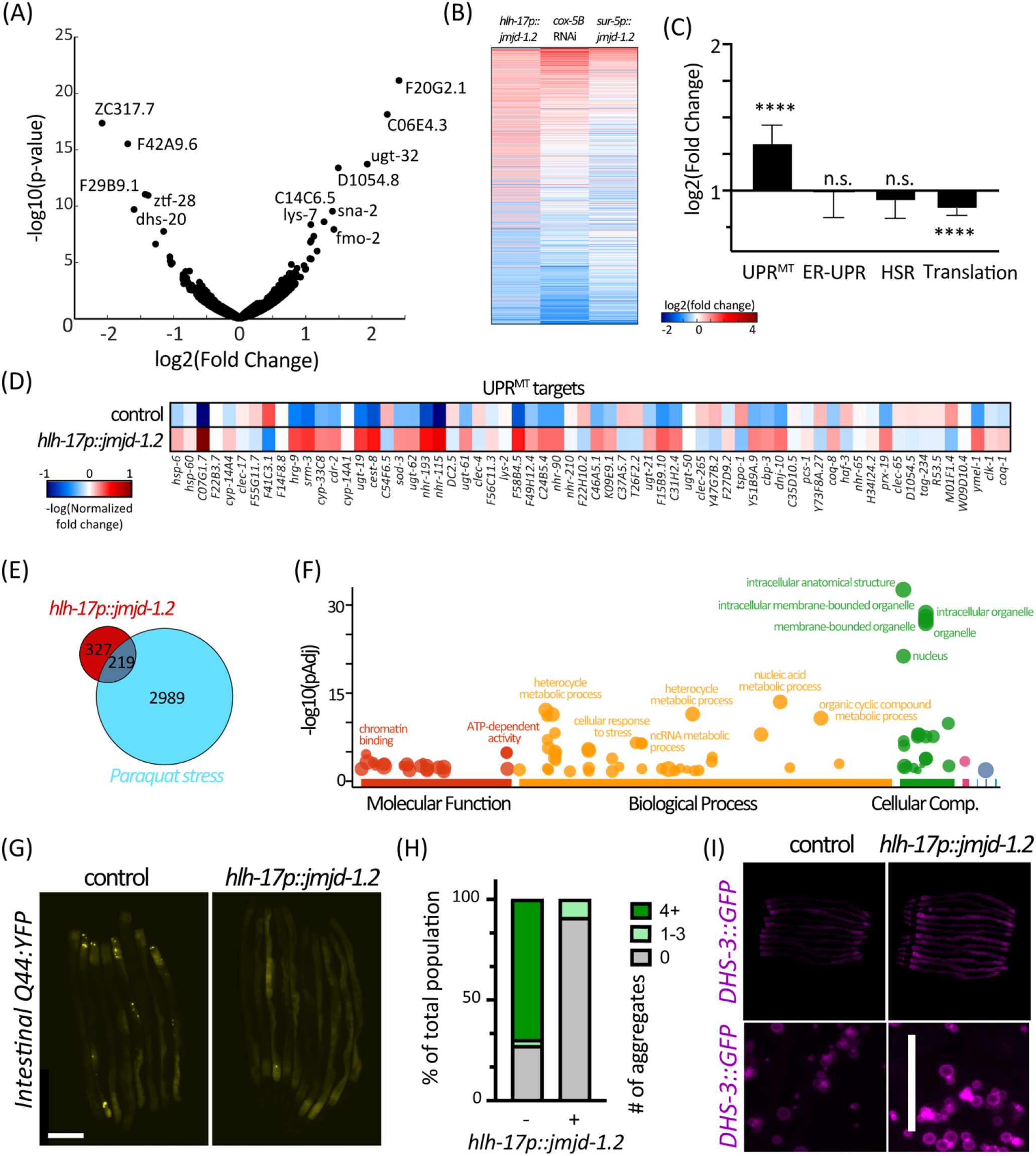
Activation of *jmjd-1.2* in CEPsh glia reduces protein aggregation, increases lipid content in the intestine, and triggers the UPR^MT^ transcriptional program. (A) Changes in gene expression in CEPsh glia expressing *jmjd-1.2*, as compared to N2 control animals, are represented in a volcano plot. (B) Differentially expressed genes in glial *jmjd-1.2* worms compared to their change in published datasets of mitochondrial stress induced by ETC knockdown by RNAi (*cox-5B* RNAi) or over-expression of *jmjd-1.2* in all worm tissues (*17*). (C) The average change in gene expression of the indicated gene groups (see methods), as compared to N2 control animals. UPR^MT^ genes are shown in (D). (E) Overlap of induced genes in glial *jmjd-1.2* worms with genes induced in paraquat stress. (F) Gene enrichment analysis was plotted using gProfiler (*45*). (G) Representative fluorescent micrographs of protein aggregation in the intestine of worms expressing *jmjd-1.2* under the CEPsh glia (*hlh-17)* promoter. Scale bar, 250 μm. (H) Quantification of number of aggregates per worm using Fiji local extrema analysis (n<30). (I) Representative fluorescent images of lipid droplets reporter animals (*DHS-3::GFP*), either in control animals or animals expressing *jmjd-1.2* in CEPsh glia cells. Significance was assessed relative to control by unpaired Student’s t test. n.s. not significant, ****P < 0.0001.

Further enrichment analysis of the upregulated genes for different gene groups revealed an overrepresentation of genes involved in organellar processes, chromatin-related pathways, metabolic processes, and stress pathways (**Fig. 3F**). To validate a physiological outcome of the increase in genes involved in stress response and protein homeostasis, we measured the capacity of animals to clear aggregation-prone proteins. Specifically, we measured the aggregation of a polyglutamine (polyQ) tract in intestinal cells (*27*), which showed a reduction in aggregation upon UPR^MT^ activation in glia (**Fig. 3, G and H and Fig. S3A**). In addition, our gene expression analysis suggested that there is also a rewiring of metabolic pathways, including lipid-related genes, which we validated directly by observing an increase in lipid droplet levels and lipid content in the intestine (**Fig. 5I and fig. S3 B to E**). These data, collectively with intestinal imaging of *hps-6p::GFP,* polyQ, and lipid content suggests that glial *jmjd-1.2* triggers a transcriptional program across the animal, which impacts UPR^MT^, protein homeostasis, and lipid levels..

### Glia utilize small clear vesicles to communicate with other tissues

The coordination of a whole animal response upon UPR^MT^ activation in CEPsh glia likely requires the transmission of a specific signal(s). The transfer of information between neurons and from neurons relies on distinct secretion pathways, with different encapsulated cargo. Previously, we found that activating the UPR^MT^ in neurons can trigger the UPR^MT^ in the intestine, utilizing the secretion of dense core vesicles (DCVs) (*8*, *10*, *11*), a pathway utilized for the secretion of polypeptide hormones and neuropeptides. These are synthesized as precursors and packaged into dense-core vesicles, processed, and depend on the Ca2+-dependent activator UNC-31/CAPS protein for secretion (*28–30*). In contrast, neurotransmitters, which are generally packaged in small clear vesicles (SCVs), and their release is dependent UNC-13/Munc13 (*31*), were not found to be involved in the activation of UPR^MT^ by neurons (*10*).

We examined whether the cell non-autonomous activation from CEPsh glia depends on the exocytosis of DCVs and/or SCVs, by mutating components important for their exocytosis, *unc-31* and *unc-13*, respectively. We found that the activation of UPR^MT^ in intestinal cells depends on both functional UNC-13 and UNC-31, and on neuropeptide processing via EGL-3 as visualized by the loss of intestinal *hsp-6p::GFP* induction in these mutants (**Fig. 4, A and B**). We also tested for another known signaling molecule required for non-autonomous communication of mitochondrial stress from neurons, the WNT signal EGL-20/WNT5A (*11*). Similar to neuronal signaling, we find that *egl-20* is also required for the distal activation of *hsp-6p::GFP* initiated from glia (**fig. S4**). Taken together, our results indicate that glial *jmjd-1.2* activation shares similar genetic requirements in secretion pathways with that of neuronal UPR^MT^ activation with the additional requirement of SCVs (*unc-13*). The requirement of SCVs is perhaps the most surprising since it clearly differentiates glial signaling from neuronal signaling, as SCVs were previously shown to be dispensable for the non-autonomous communication from neurons. We therefore hypothesize that glia communicate to neurons via *unc-13* (SCVs) and neurons communicate to the periphery via *unc-31* (DCVs) signaling.

**Figure 4:**
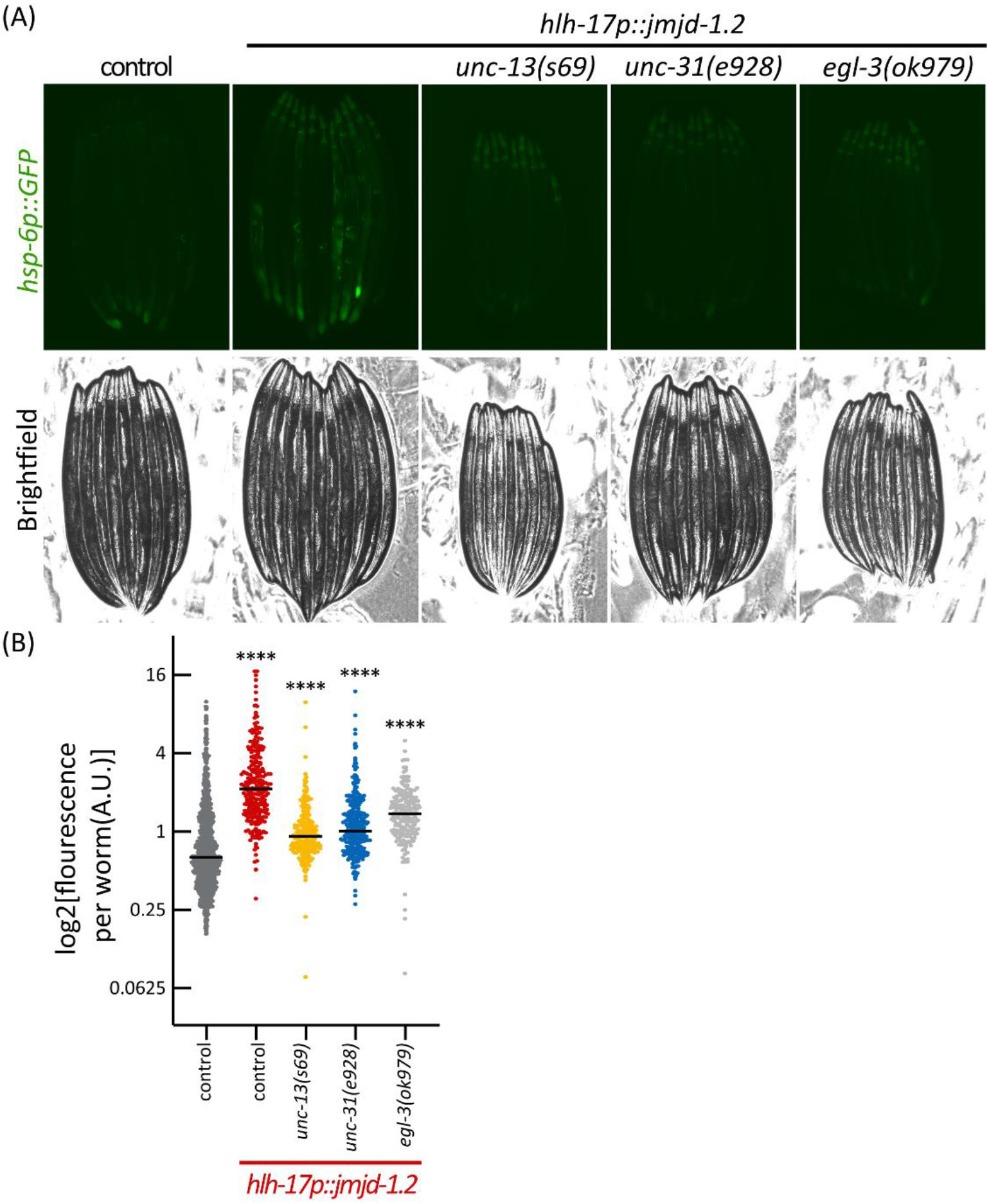
Cell non-autonomous activation of the UPR^MT^ in the intestine depends on the secretion of SCVs, DCVs, and neuropeptide processing. (A) Representative fluorescent micrographs of UPR^MT^ reporter worms (*hsp-6::GFP*) for animals with mutations in the secretion of small-clear vesicles (SCV, *unc-13*), dense-core vesicles (DCV, *unc-31*), and neuropeptide processing (*egl-3*). Scale bar, *25*0 um. (B) Quantification of UPR^MT^ reporter worms (*hsp-6::GFP*) as in Figure 1E (n>200).

To directly test this hypothesis and uncover in which tissues (glia vs neurons) *unc-13* and *unc-31* participate in the UPR^MT^ signaling, we knocked out *unc-13* and *unc-31* in a tissue-specific manner using the FLP/FRT system (*32*). This system allows deleting a portion of a gene by expressing the flipase, FLP D5, in specific tissues of interest (CEPsh glia vs pan-neuronal) in combination with an allele of the gene of interest (*unc-13* or *unc-31*) to which two copies of the FRT sequence have been introduced using CRISPR/Cas9 genome editing (**Fig. 5A and fig. S5, A to C**). The resulting animals have spatially distinct genotypes, with a defect in the secretion of SCVs or DCVs exclusively in one tissue. Strikingly, we observed that knocking out *unc-13* specifically in glia, or *unc-31* specifically in neurons, was able to alleviate the activation of the UPR^MT^ in the intestine (**Fig. 5, B to E**). Moreover, rescuing *unc-13* specifically in neurons, under the *snb-1* promoter, was not able to restore the activation of UPR^MT^ in the periphery (**fig. S5D**). These results, together with our previous work on neuronal activation of the UPR^MT^, indicate a linear model, in which glia secrete a molecule by SCVs (*unc-13*), that is perceived by neurons, which then relay the information to the periphery via DCVs (*unc-31*).

**Figure 5:**
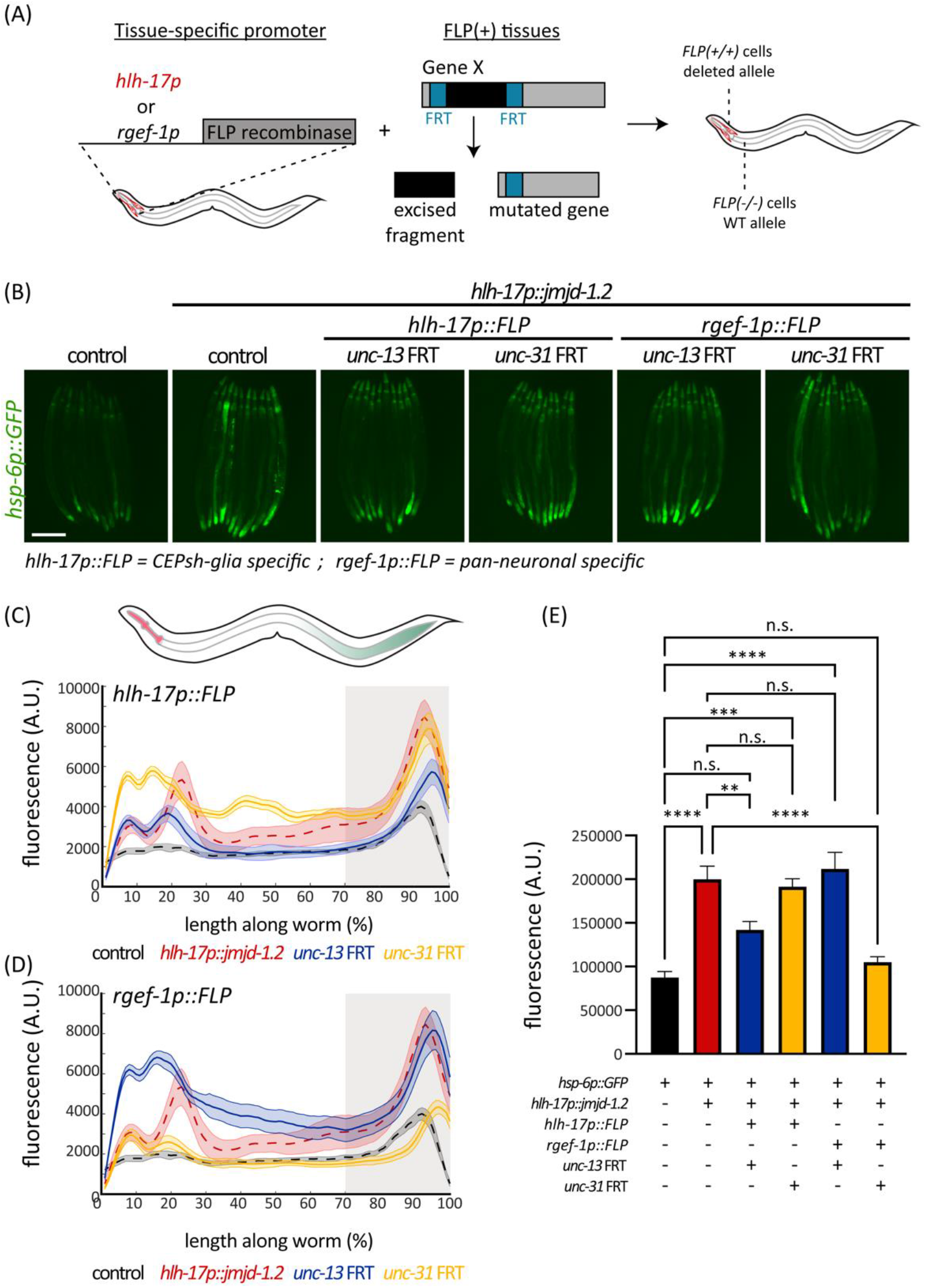
CEPsh glia require functional UNC-13 (SCV), while neurons require UNC-31 (DCV), to activate UPR^MT^ in the periphery. (A) Schematic of spatial mutation strategy. Expression of the flipase FLP D5 under the CEPsh glial-specific (*hlh-17p*) or pan-neuronal (*rgef-1p*) promoters, in combination with an independent FRT (FLP recognition target) allele of a gene of interest results in a tissue-specific mutation. (B) Representative fluorescent micrographs of the indicated strains at day 1 adult animals. Scale bar, *25*0 um. (C & D) Median spatial profiles of the indicated animals (see methods), for depletion in CEPsh glial cells (C) or neuronal cells (D) quantified with large-particle biosorter (n>50). The integral of the 30% most posterior portion of the animals was calculated and plotted in (E). One-way analysis of variance (ANOVA) Tukey’s multiple comparisons test. n.s. not significant, ***P < 0.001, ****P < 0.0001.

### Activation of UPR^MT^ in glia improves neuronal protein homeostasis

Considering our model whereby activation of glial UPR^MT^ results in a glia-to-neuron signal, we next questioned whether glial UPR^MT^ is able to drive improved protein homeostasis in neurons. Indeed, our data and previous studies (*10*) showed that non-autonomous UPR^MT^ signals can promote protein homeostasis in the target tissues. Moreover, the importance of protein homeostasis in neurons is best illustrated by neurodegenerative diseases, in which protein aggregation occurs (*5*), of which glial health has been suggested as a critical factor in maintenance of neuronal fitness (*14*). Thus, we employed the well-established Huntington’s disease model in *C. elegans*, in which a 40-repeat polyglutamine tract (Q40) is expressed in all neuronal cells. Interestingly, Q40 is able to bind directly to mitochondria, and has been shown to also affect mitochondria directly in the same tissue (*10*, *33*). We activated the UPR^MT^ in glia and measured a battery of phenotypes associated with the neuronal HD model. Notably, activating UPR^MT^ in glia was able to rescue the thrashing and chemotaxis defects observed in neuronal Q40 animals (**Fig. 6, A and B and fig. S6**).

**Figure 6:**
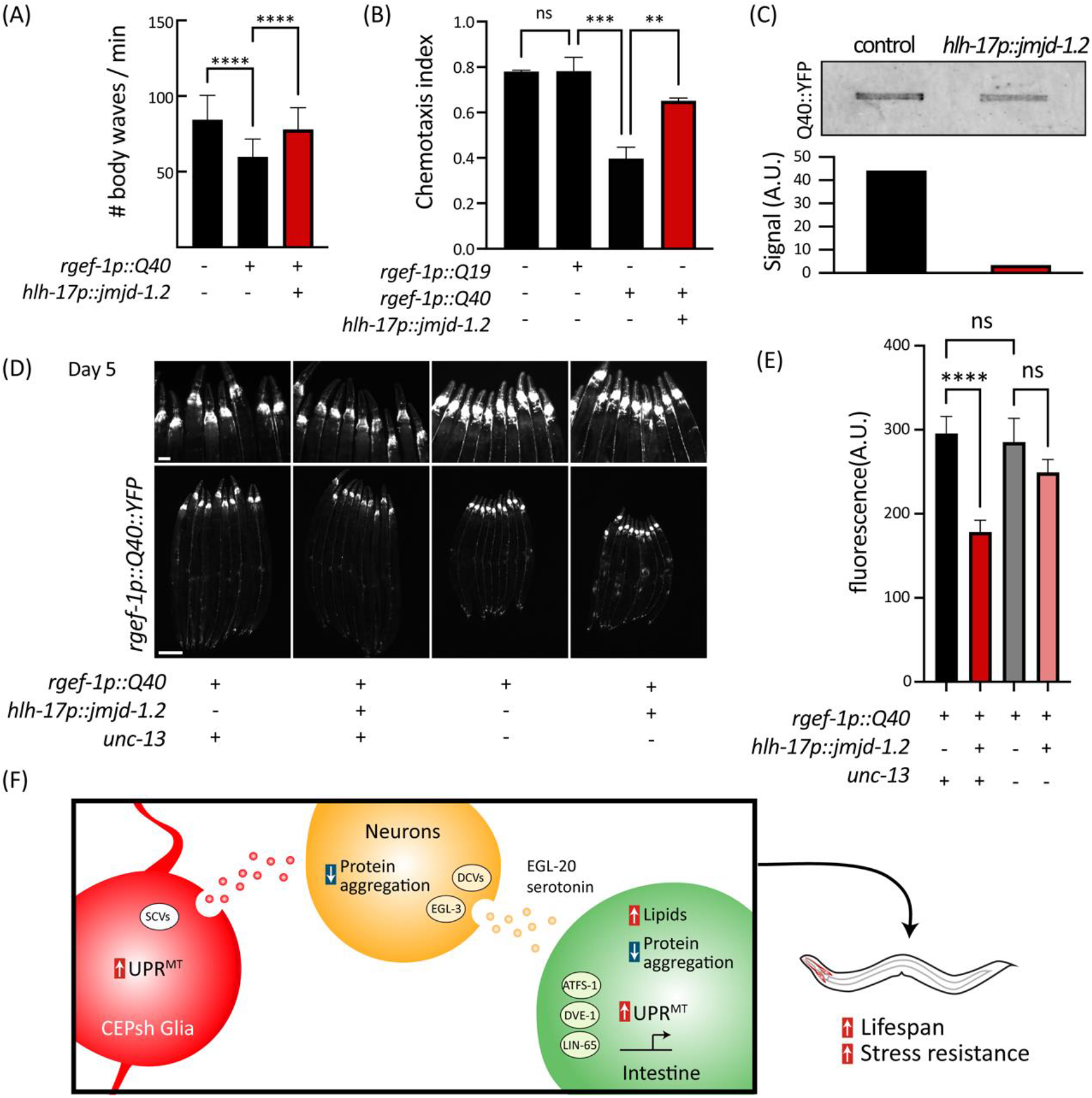
Glial *jmjd-1.2* rescues protein aggregation in neurons in a Huntington’s disease (HD) model via SCVs. (A) Thrashing of animals expressing the aggregating polyglutamine tract Q40, with and without glial *jmjd-1.2*, measured using WormTracker (n>100) (B) Chemotaxis index of worms towards benzaldehyde (n>200). (C) Filter Retardation Assay for Q40::YFP, with and without *glial jmjd-1.2* (top) and its quantification (bottom). (D) Representative fluorescent micrographs of Q40::YFP for the annotated genotypes, on D5 of adulthood. See Figure S6 for images of worms at D1. (E) Quantification of Q40 signal in the head region of the animals (n>30) using Fiji. One-way analysis of variance (ANOVA) Tukey’s multiple comparisons test. n.s. not significant, *P < 0.05, **P < 0.01, ***P < 0.001, ****P < 0.0001. (F) Model of communication from glial cells to peripheral tissues. CEPsh glia utilize SCVs upon UPR^MT^ activation to signal to neurons, which reduce protein aggregation and utilize DCVs, neuropeptide processing, and a WNT-ligand to drive protein homeostasis and metabolic changes in the periphery.

The polyglutamine model is also useful for directly measuring protein aggregation, as the Q40 tract is tagged with a YFP. We imaged worms and observed decreased aggregation of the protein in late age (**Fig. 6, D and E and fig. S6C**). We further verified the aggregation using a filter-trap assay (*34*), where we observed less aggregation when activating the UPR^MT^ in glia (**Fig. 6C**). Next, we tested whether these effects are dependent on SCVs. Indeed, intact UNC-13 signaling was required for alleviating the aggregation in neurons (**Fig. 6, D and E**). Finally, we asked whether this protection by glia is a general attribute of CEPsh glia, by activating other protein homeostasis pathways in CEPsh glia and testing their ability to improve protein homeostasis in neurons. To this end, we employed our previous model of activating the UPR^ER^ in CEPsh glia, using an overexpression of the spliced version of the transcription factor *xbp-1s* (*16*). Interestingly, we were not able to observe any beneficial effects (**Fig. S6D**), highlighting that the ability of glia to improve neuronal protein homeostasis is unique to the UPR^MT^.

## DISCUSSION

In this study, we discovered a role for a small subset of glial cells in sensing mitochondrial stress and signaling a beneficial cell-non-autonomous communication to neurons. This signal drives a coordinated change in mitochondrial protein homeostasis across tissues, via a relay-cellular pathway mediated by neurons. We found that communication from glia occurs as a two-step process: the astrocyte-like CEPsh glial cells secrete a signal via SCVs, and subsequently, neurons relay the signal to peripheral tissues by utilizing DCVs, and the WNT ligand EGL-20 (**Fig. 6F**).

We found that different subpopulations of glial cells can regulate longevity and coordinate the communication of mitochondrial proteotoxic stress across the tissues of the animal. We were most struck by the CEPsh glia, a cell type that includes only four individual cells, that resulted in multiple physiological benefits, including increased lifespan and stress resistance, with full dependency on the functionality of the regulators involved in the UPR^MT^. Our efforts to uncover and disentangle the requirement of genes involved in intercellular signaling revealed that CEPsh glia utilize small clear vesicles to signal the mitochondrial proteotoxic stress. Indeed, mammalian astrocytes were suggested to possibly express Munc13, in cultured and freshly isolated astrocytes (*35*, *36*), as similarly observed in RNA-seq data from isolated CEPsh glia in nematodes (**Fig. S5E**) (*37*), and our findings may provide the context in which these vesicles are utilized.

Our dissection of the functional importance of different genes in communicating the UPR^MT^ from glia to the intestine suggests that the signal is mediated by neurons. We were thus inspired to test whether the internal state of neurons themselves is altered upon receiving the UPR^MT^ from glia. Interestingly, we found that glial UPR^MT^ can promote protein homeostasis in neurons and protect them from the detrimental effects of polyQ expression. Conventionally, neurons have been at the center of studying neurodegenerative diseases and glial cells, such as astrocytes, have been implicated in neurogenerative diseases, mostly by ablation experiments showing that glial cells can exacerbate disease pathology (*13*). Thus, whether the loss of glial cells directly caused disease phenotypes was not understood, and most studies concluded that the breakdown of neuronal function was the true culprit. However, our study supports a redefining role of astrocytes from simply supporting neuronal cells to playing an active role in sensing stress and communicating a beneficial signal to neurons, which can then mediate a whole-organism response to stress. These findings have potentially huge ramifications for the study of neurodegenerative diseases, as it shifts attention to glial cells and asks the question of whether glial cells should be the true target for potential therapies in neurodegenerative disorders. Indeed, increased activity of astrocytes identified by those with higher levels of the glutamate transporter, GLT-1, was found to preserve cognitive function in patients with Alzheimer’s (*38*).

An important question raised by our study is whether CEPsh glia can uniquely sense mitochondrial stress. Our data show that activation of the UPR^MT^, but not the UPR^ER^, in CEPsh glia resulted in increased neuronal protein homeostasis. There are two plausible explanations for this: first, it is possible that a distinct glial population is required for sensing and communicating ER stress. Indeed, other studies have shown that neuronal cells differ in their capacity to sense and signal distinct types of UPR^ER^ signals (*39*), and it is entirely possible that subsets of glia similarly have distinct abilities to sense and signal stress. Another possibility is that mitochondrial stress signals are uniquely able to activate protein homeostasis machinery adept at clearing polyQ aggregates, whereas other stress signals may activate protein homeostasis machinery for other forms of protein dysfunction. Indeed, previous studies have shown that polyQ aggregates can bind mitochondria and induce the UPR^MT^ (*10*), suggesting there are organelle-specific effects of certain classes of aggregates.

Finally, we found that glia have the ability to improve protein homeostasis not only on the physiological level (thrashing, chemotaxis) but also on the molecular level, measuring the aggregating protein itself. Similarly, recent work in mice found that indeed astrocytes, through activation of the JAK2-STAT3 pathway, are able to clear protein aggregates in neurons in a mutant huntingtin mouse model (*40*), highlighting our findings of a UNC-13-related improvement of protein homeostasis in the glial-neuron axis. Of note, UNC-13/Munc13 has previously been linked also to another neurodegenerative disease, Amyotrophic Lateral Sclerosis (ALS). Genome-wide association studies linked UNC13A to ALS as mutations associated with higher susceptibility and shorter survival (*41–43*) in individuals with ALS, a link that is evolutionarily conserved in *C. elegans* (*44*). Whether the involvement of UNC-13 in ALS stems from its glial functions remains to be explored.

In summary, our work not only highlights the importance of glial cells in communicating mitochondrial proteotoxic stress, and coordinating between tissues, but also mechanistically mapped SCVs as the secretion pathway utilized from these cells. Our study places CEPsh glia as upstream regulators of coordination of stress responses and underlines their role as a major signaling hub that can affect cellular and organismal homeostasis. In combination with our findings on protein aggregation in neurons, our results underscore the importance on examining the roles and mechanisms used by glial cells to regulate protein homeostasis within the nervous system and in the periphery. Advancing our knowledge of how glial cells regulate protein homeostasis in the context of mitochondrial stress will be critical in ongoing efforts to understand the glia-neuron communication axis, in neurodegenerative diseases, and in aging.

## Acknowledgments

We are grateful to Larry Joe, Melissa Sanchez, Dr. Tslil Ast, Dr. Naama Aviram, Dr. Ranit Kedmi, and all members of the Dillin lab for technical support and sharing of reagents and equipment.

## Author contributions

R.B.Z: Conceptualization, Methodology, Software, Validation, Formal analysis, Investigation, Resources, Data Curation, Writing - Original Draft, Writing - Review & Editing, Visualization, Supervision, Project administration. **E.S.**: Methodology, Investigation, Resources. **H.R.H.**: Conceptualization, Methodology, Investigation, Formal analysis. **J.D.**: Methodology, Investigation, Validation. **S.U.T.**: Methodology, Investigation, Validation, Formal analysis. **Q.A**: Methodology, Investigation. **T.B.** Methodology, Investigation. **J.P.** Methodology, Investigation. **J.G.D.**: Methodology. **R.H.S.**: Methodology, Validation, Formal analysis, Investigation, Resources, Writing - Original Draft, Writing - Review & Editing. **A.D.**: Conceptualization, Writing - Review & Editing, Supervision, Project administration, Funding.

## Data and materials availability

All data required to evaluate the conclusions in this manuscript are available within the manuscript and Supplementary Materials. All strains synthesized in this manuscript are derivatives of N2 or other strains from CGC and are either made available on CGC or available upon request to the lead contact. Plasmids cloned to generate the used strains are available upon request. RNA-seq datasets supporting the conclusions of this article are available in the NCBI Sequence Read Archives repository, accession number BioProject PRJNA858426.

## Methods

### Strains and maintenance

All *C. elegans* strains are derivatives of the Bristol N2 wild-type strain from the Caenorhabditis Genetics Center (CGC) and are listed in Table 1. All worms were grown at 15° to 20°C on NGM (nematode growth media) agar plates, fed with OP50 *E. coli* B strain as a food source, for general maintenance, and handled as previously described (*50*). For all experiments, worms were switched to HT115 *E. coli* K12 strain after synchronization using bleaching. Worms were grown for at least 3 generations at 20°C on OP50 prior to synchronization to acclimate to the temperature. HT115 bacteria were carrying the pL4440 empty vector control or expressing double-stranded RNA containing the sequence against a target gene, as specified in the text. RNAi clones against *atfs-1* and *daf-2* were acquired from the Vidal RNAi library (*51*) and verified using Sanger Sequencing. All experiments were performed on age-matched animals synchronized using a standard bleaching protocol, to Day 1 of adulthood, or aged to later ages, as specified. Synchronization was achieved by washing animals fed with OP50 *E. coli* with M9 solution (22 mM KH_2_PO_4_ monobasic, 42.3 mM Na_2_HPO_4_, 85.6 mM NaCl, and 1 mM MgSO_4_), bleached using a solution of 1.8% sodium hypochlorite and 0.375 M KOH diluted in DDW, for 4-5 minutes, until all carcasses were digested. Intact eggs were then washed 4× with M9 solution, and intact eggs were verified under the microscope after seeding.

**Table 1:**
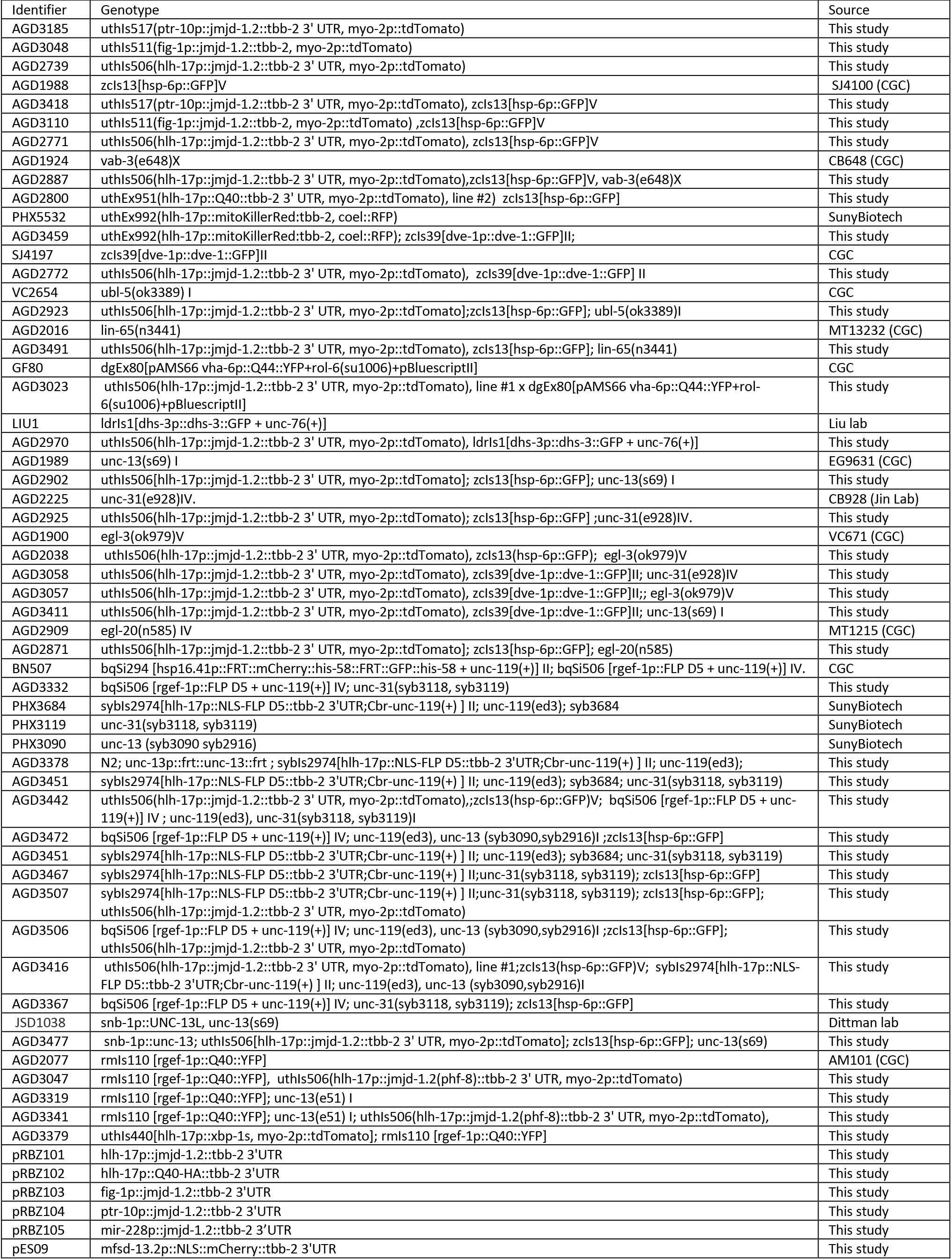
Strains and plasmids used in this study.

Strains were generated by cloning the cDNA of interest under the relevant promoter, using Gibson Assembly (*52*). The cDNAs of *jmjd-1.2* and the polyglutamine Q40 were amplified from pCM407 (*17*) and pJKD101 (*10*), respectively. The promoters *ptr-10p, fig-1p*, and *hlh-17p* were sub-cloned from pAF12, pAF5 and pAF6 (*16*), respectively. Synthetic DNA of *tbb-2* 3’UTR and *mfsd-13.2* promoter was synthesized by Twist Bioscience. Wild type N2 strain worms were injected with the construct of interest, along with a *myo-2p::tdtomato or unc-122p::RFP* co-injection marker. Worms were selected under a fluorescent microscope and then integrated by UV irradiation. Integrated lines were backcrossed 8 times to a wild-type N2 strain. For same-orientation FRT insertions; *unc-13* between exon 18 and 19 and between exon 20 and 21, and for *unc-31*, insertions between exon 1 and 2, and 2 and 3 were made using CRISPR/Cas9 by SUNY biotech, and successful cutting was verified using PCR and gel electrophoresis, followed by sequencing (see Figure S5A).

### Microscopy and of UPR^MT^ reporters

Imaging of *hsp-6p::GFP* and DVE-1::GFP fluorescent reporters was done as previously described (*53*). Briefly, animals were grown on standard RNAi plates from hatch at 20°C until day 1 of adulthood. Animals were picked under a standard dissection microscope with white light at random to avoid biased sampling. Then, animals were anesthetized using a 10-15ul drop of 100 mM sodium azide (NaN3) on NGM plates with no bacteria and aligned to have the same orientation. Images were captured on a Leica M250FA stereoscope equipped with a Hamamatsu ORCA-ER camera driven by LAS-X software, or using an Echo Revolve R4 microscope equipped with an Olympus 4x Plan Fluorite NA 0.13 objective lens, a standard Olympus FITC filter (ex 470/40; em 525/50; DM 560). Brightfield and fluorescent images were acquired, with exposure time and laser intensity matched within experiments. Each micrograph contained 10 individual worms and was independently replicated at least three times.

### Large-particle flow cytometry

To quantify fluorescent reporters, flow cytometry using a Union Biometrica bioSorter (cat #250-5000-000) was done as previously described (*53*). Briefly, staged worms were washed off plates using M9, allowed to settle by gravity, and washed once with M9 to separate from eggs. The signal was collected for time of flight (length) and extinction (thickness) of animals, along with the GFP and RFP. Data were collected gating for size (time of flight [TOF] and extinction) to exclude eggs. Data are represented as an integrated intensity of fluorescence normalized to the size of the animal using the integrated GFP output and dividing by the extinction and time of flight. All data that exceed the measurement capacity of the PMT, calculated as a signal of 65355, are considered saturated and are censored from the calculation. For spatial profiles, the complete profiles were extracted, and worms were aligned according to their *myo-2p::tdtomato* (red head) signal using MATLAB (MathWorks), and binned into 100 bins to account for differences in animal length (n>50). Then, the average profile and SEM was calculated on binned profiles. For *hsp-6p::GFP* worms, which do not harbor a *myo-2p::tdtomato* co-injection marker, the profiles were aligned according to the GFP signal, with the highest peak of signal defined as the posterior intestine.

### Lifespan analysis

Lifespan analyses were performed as previously described (*53*). All animals were grown at 20°C on HT115 *E. coli*, with 80-120 animals used per condition and scored every day or every other day, starting from day 1 of adulthood. Animals were moved away from progeny onto fresh plates for the first 5-7 days until progeny were no longer visible. Animals with bagging, vulval explosions, or other age-unrelated deaths were censored and removed from quantification. At least two independent replicates per condition. Analyses were done using Prism 8 (GraphPad). *P*-values were calculated using the log-rank (Mantel–Cox) method.

### Paraquat resistance assay

Resistance to oxidative stress generated by exposure to 100mM paraquat (Sigma-Aldrich 36541) (*19*) was done as previously described (*53*), with three biological replicates per condition. Briefly, fresh paraquat was prepared in M9 solution, and aliquoted into a flat-bottom 96-well plate, with 5 animals per well, with 12 wells per strain (n=60). Every two hours, animals were scored for death in each well. For all paraquat assays, *daf-2* RNAi is used as a positive control as *daf-2* animals exhibit significant resistance to oxidative stress using this assay (*53*).

### RNA sequencing and analysis

Animals were synchronized using a standard bleaching protocol and all RNA collection was performed at day 1 of adulthood fed with HT115 bacteria. A total of 1000-2000 Day 1 animals were harvested using M9, and animals were pelleted by centrifugation. M9 was subsequently aspirated and replaced with TRIzol solution. Worms were freeze-thawed 3× with liquid nitrogen, and a ∼30-s vortexing was performed before each refreeze. After the final thaw, chloroform was added at a 1:5 ratio (chloroform:TRIzol), and aqueous separation of RNA was performed via centrifugation in a heavy gel phase-lock tube (VWR, 10847--802). The aqueous phase was collected, mixed with isopropanol at a 1:1 ratio, and then RNA purification was performed using a QIAGEN RNeasy Kit as per the manufacturer’s directions. Library preparation was performed using Kapa Biosystems mRNA Hyper Prep Kit. Sequencing was performed at the Vincent J.

Coates Genomic Sequencing Core at the University of California, Berkeley using an Illumina NovaSeq SP SR100. For N2 control and *hlh-17p::jmjd-1.2*, two or three biological replicates were done, respectively. Reads were aligned using HISAT2 (Version 2.2.1) (*54*) with WBcel235 as the reference genome, quantified using featureCounts (*55*), and DEG were calculated using DESeq2 (*56*). Gene groups for analyses: mitochondrial UPR were defined as previously annotated by Soo et al. (*57*), ER-UPR (GO:0030968), HSR (GO:0009408), and translation (GO:0006412) genes were defined using Gene Ontology (*58*, *59*). Comparison to other gene expression collections was done using WormExp (*60*). Gene enrichment was done using gProfiler (*61*) and GOrilla (*62*), for significantly changing genes (P<0.05). Plots were generated using MATLAB (MathWorks).

### BODIPY staining

Neutral lipid measurements was done using BODIPY 409/503 staining as previously described (*32*), with three biological replicates. Briefly, worms were synchronized by bleaching, grown on EV, and harvested at L4. Worms were washed three times in M9 buffer. Then 4% PFA was added and incubated for 15 min to fixate the samples. Next, the sample was frozen in liquid nitrogen, and immediately thawed (one time), followed by three washes in 1X PBS. Staining was carried out by incubating worms in 500ul of 1ug/ml BODIPY 493/503 for 1 HR at RT shaking, in the dark. After incubation, worms were washed with M9 three times to remove BODIPY. Worms were then left O/N at 4°C, in shaking, in 1X PBS. Finally, worms were imaged using a fluorescent microscope, or sorted using COPASS BioSort (see relevant section).

### Q40 aggregation

Worms were bleached, and eggs grown to day 1 of adulthood on HT115 bacteria. Worms were imaged at Day 1 and Day 5 of adulthood, and then imaged using fluorescent microscopy (see relevant section). Q40::YFP was expressed in neurons or intestine using promoters *rgef-1p* or *vha-6p* respectively. For normalization of protein, standard western blotting protocols were used. 100 age-synchronized animals were hand-picked into an M9 solution and lysed in SDS-lysis buffer using 3x cycles of 100 °C boiling for 10 min followed by flash-freezing in liquid nitrogen. The sample was then centrifuged at 10,000 x g for 5 min to pellet cell debris. The top 15 µL was pulled off and run on a 4-20% Criterion TGX Precast gel using a standard protocol and probed using an anti-GFP antibody, fluorescent LI-COR secondary, and imaged on a LICOR Odyssey M similar to filter trap assay below.

For intestinal polyQ imaging, osmotic stress was applied to worms at day 1 of adulthood by moving animals on to a plate with 500 mM NaCl and imaged after 4 hours of incubation. Three biological replicates were done.

### Thrashing assay

Thrashing of worms was measured using the WormLab (MBF Bioscience) system. Worms were grown to day 1 of adulthood, washed off plates using M9 buffer, and then allowed to settle by gravity in an Eppendorf tube, washed once, and then moved to an empty 6mm plate (without NGM). Worms were video recorded, in three biological replicates, and their thrashing (body waves) were measured using WormLab 2.0 software. The size of each worm (length and width output from WormLab) to gate for worm events, and thrashing was averaged across the gated population.

### Chemotaxis assay

Animals were synchronized by bleaching, and assay was conducted at day 1 of adulthood as previously described. Briefly, animals were raised on EV bacteria until day 1 of adulthood and washed three times using M9 buffer by gravity settling. Chemotaxis assay plates were prepared: 100mm NGM plates, with 1uL attractant (Benzaldehyde 1:200, Diacetyl 1:1000) and 1uL diluent (EtOH) on opposite sides, with 1uL of sodium azide in the same location, to immobilize worms once reaching their target. In equal distance, 30-50 worms were dispended and scored after 60 minutes. Chemotaxis index was calculated by I = (# worms at attractant at 60’ - # worms at diluent at 60’) / total number of worms. Assays were repeated at least 3 times, and significance was assessed by one-way ANOVA with Tukey’s multiple comparisons test.

### Filter trap retardation assay

Worms grown to day 1 of adulthood were washed off the plate using M9, harvested and flash-frozen in liquid nitrogen. Then, animals were suspended in lysis buffer (100 mM Hepes pH 7.4, 300 mM NaCl, 2 mM EDTA, 2% Triton X-100, with EDTA-free protease inhibitor cocktail (Roche). Next, animals were homogenized using a Precellys Tissue Homogenizer, with glass and zirconium beads (2mm). Lysates were centrifuged (8000g for 5 minutes at 4 °C), supernatant moved and protein quantified using BCA Protein Assay kit (ThermoFisher, 23225). Protein samples were applied on to cellulose acetate membrane with 0.22 mm pore size (Schlechtes + Schule), assembled in vacuum slot blotter (Bio-Dot, Bio-Rad). Membrane was washed with 0.2% SDS five times on the blotter and subjected to antibody incubation for detecting aggregated protein retained on the membrane. Membranes were incubated with anti-GFP antibody (1:1000 dilution in LI-COR blocking buffer) overnight in a cold room. Membrane was washed with TBST for three times, then incubated with secondary antibody (1:10000 dilution in LI-COR blocking buffer). Membranes were washed with TBST three times and imaged using Odyssey® M Imaging System (LI-COR) to visualize the protein bands. Bands were then quantified using Fiji.

## Supplementary Materials

**fig. S1-S6**

**Table S1.** DEGs in *hlh-17p::jmjd-1.2* animals.

**Table S2.** Enrichment of upregulated genes using gProfiler.

**Table S3.** GO enrichment for DEGs.

**Table S4.** Statistics for lifespan.

