## Supplemental Figures for "Glial-derived mitochondrial signals impact neuronal proteostasis and aging"

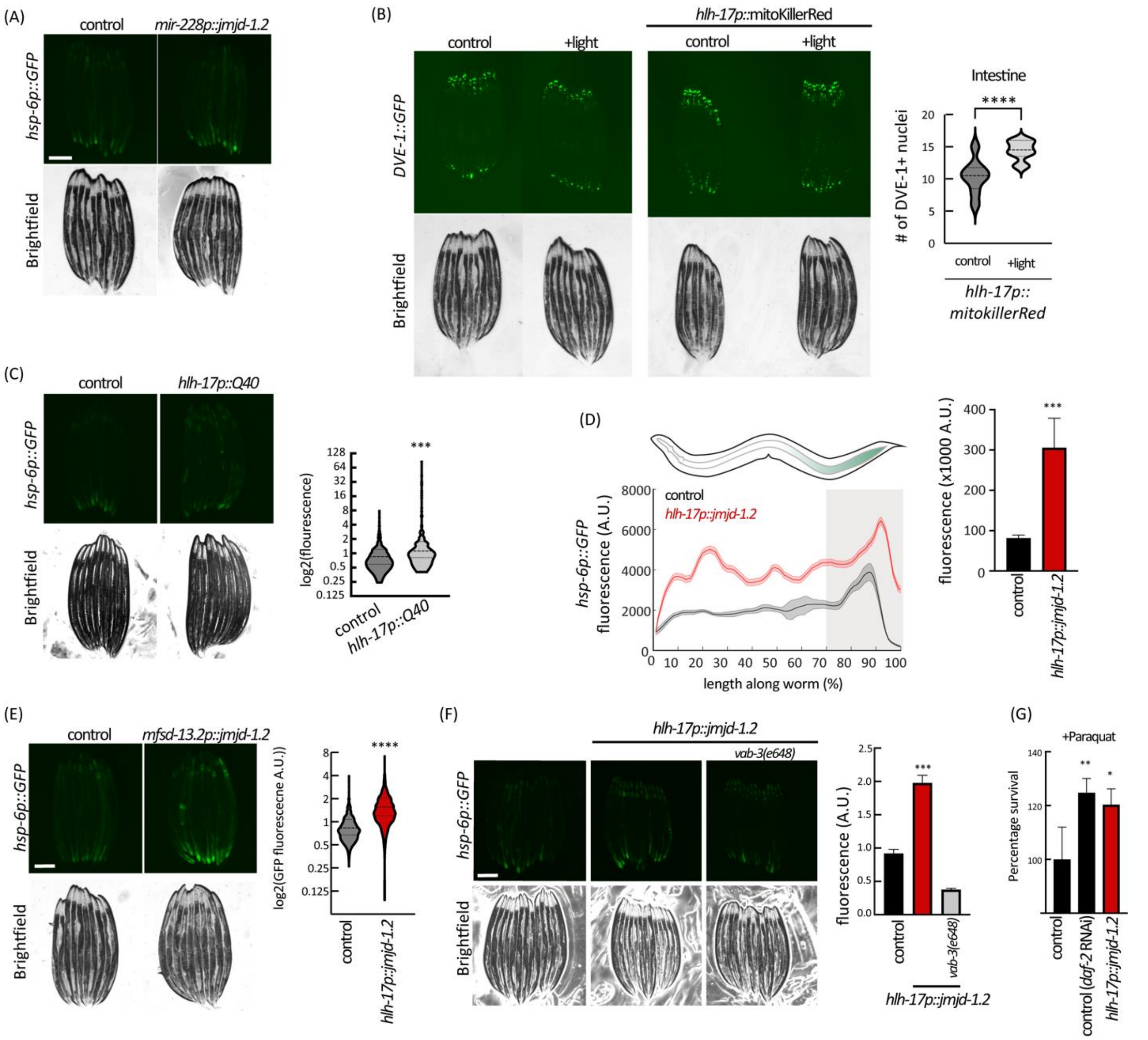


**Figure S1: Activation of *jmjd-1.2* in glia induces UPR$^{MT}$ in the periphery, related to Figure 1.**

(A) Representative fluorescent micrograph of UPR$^{MT}$ reporter *hsp-6p::GFP*, for worms expressing *jmjd-1.2* in most glial cells, under the *mir-228* promoter (Fung et al., 2020). (B) Same as S1A, for worms over-expressing mitochondrially targeted KillerRed construct, under the regulation of the *hlh-17* promoter, and the translational reporter DVE-1::GFP (left, see Figure 2A). L4 worms were irradiated with 1 min of light (543 nm excitation filter), and imaged after 24 hours, and quantified (right). unpaired Student's t test, ****P < 0.0001. (C) Same as S1A, for worms carrying an extrachromosomal array of the polyglutamine tract Q40 under the *hlh-17* promoter. (D) Worms over-expressing *jmjd-1.2* under *hlh-17p* were analyzed using a biosorter, and their spatial profiles aligned (left, see methods). The 30% most posterior part of the worm was defined as the posterior intestine, and the integral of the signal over the region was calculated (right). unpaired Student's t test, ***P < 0.001. (E) Same as S1A, for worms over-expressing *jmjd-1.2* from a promoter identified as specific for CEPsh glia (Roux et al., 2022), *mfsd-13.2*p (C27H5.4) (left), and its quantification using biosorter (right). unpaired Student's t test, ****P < 0.0001. (F) As figure S1A, for *hlh-17p::jmjd-1.2* worms in combination with a mutation in the CEPsh glia lineage mutation in *vab-3* (Yoshimura et al., 2008), and its quantification using biosorter (right). (G) Percentage survival of worm exposed to paraquat was calculated by integrating the area under the curve, and the change in resistance was normalized by the average integral of control worms (N2), set as 100%. One-way analysis of variance (ANOVA) Tukey's multiple comparisons test, $^{**}P < 0.01$, $^{***}P < 0.001$, $^{****}P < 0.0001$.

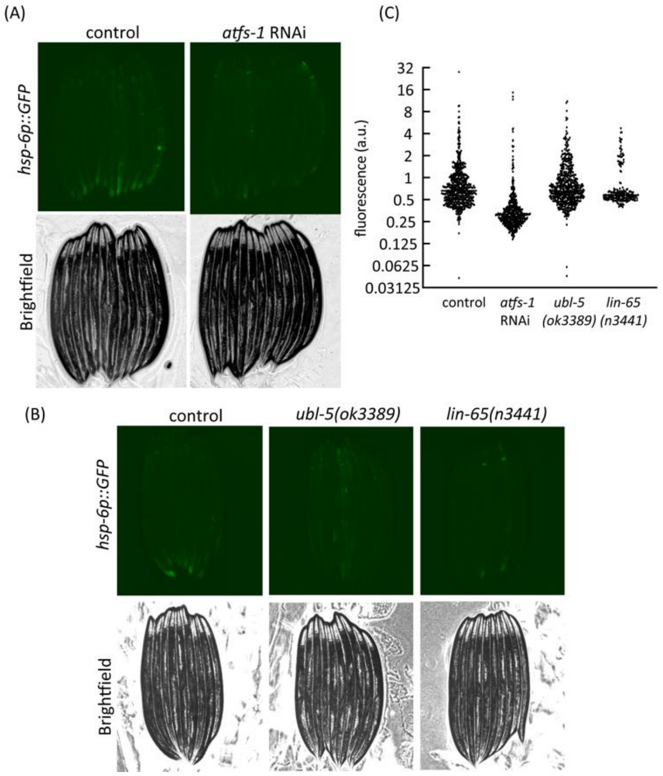


**Figure S2: Activation of UPR$^{MT}$ in the periphery depends on cell-autonomous regulators of the pathway, related to Figure 2.**

(A) Representative fluorescent micrograph of UPR$^{MT}$ reporter *hsp-6p::GFP*, for worms expressing *hsp-6p::GFP* , with RNAi against the transcription factor *atfs-1,* or with mutations against *ubl-5* and *lin-65* (B), other players in the UPR$^{MT}$ pathway. Quantified using biosorter (C).

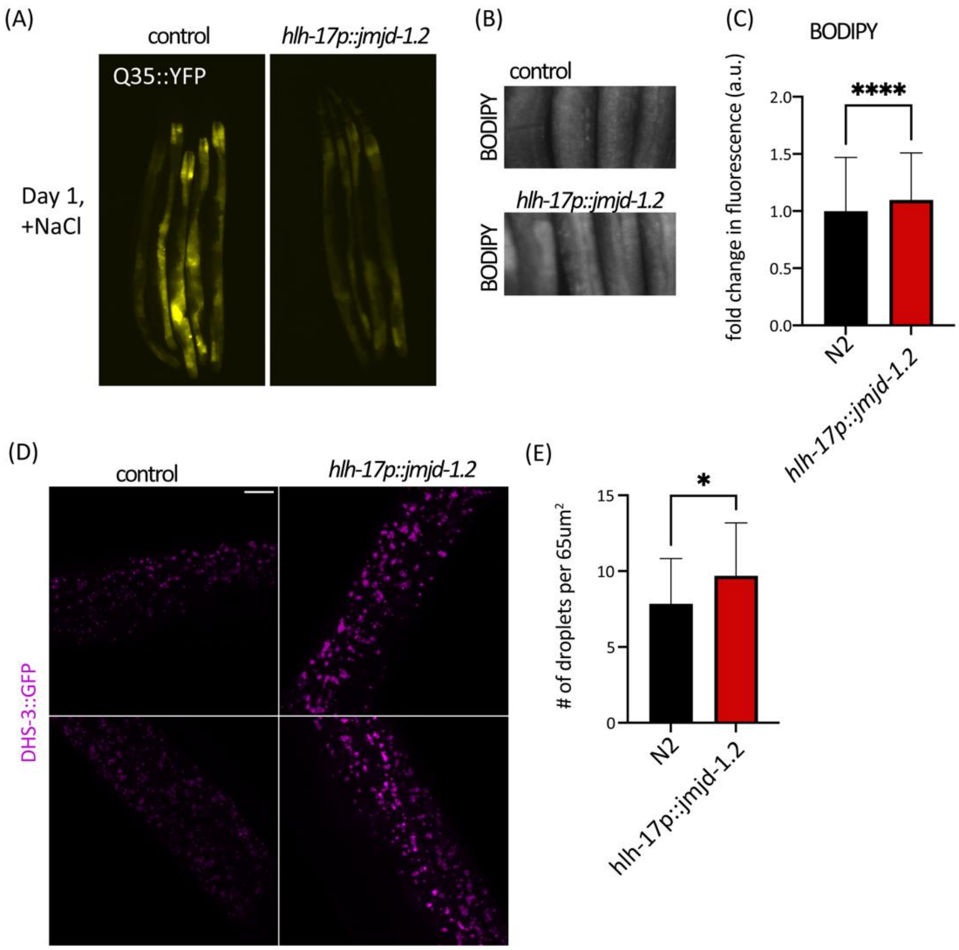


**Figure S3: Glial activation of *jmjd-1.2* improves protein homeostasis in the periphery and induces lipid droplets, related to Figure 3.**

(A) Representative fluorescent micrographs of Q35::YFP worms expressing in the intestine, after 4 hours in 400mM NaCl at D1 of adulthood. (B) BODIPY straining at D1 of adulthood and its quantification using worm biosorter (C). Imaging of DHS-3::GFP as in Figure 3 and its quantification using Fiji (E). unpaired Student's t test, $*P < 0.05$, $****P < 0.0001$.

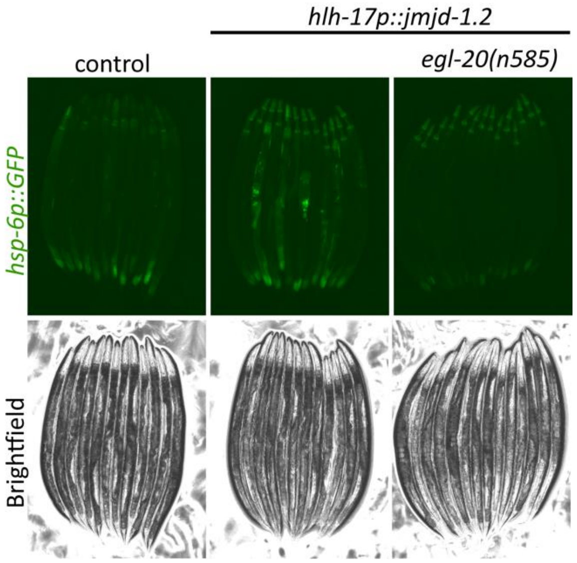


**Figure S4: Cell non-autonomous activation by glial *jmjd-1.2* depends on WNT signaling, related to Figure 4.**

Representative fluorescent micrographs as in Figure 4, for worms mutated for the WNT ligand *egl-20* (Zhang et al., 2018).

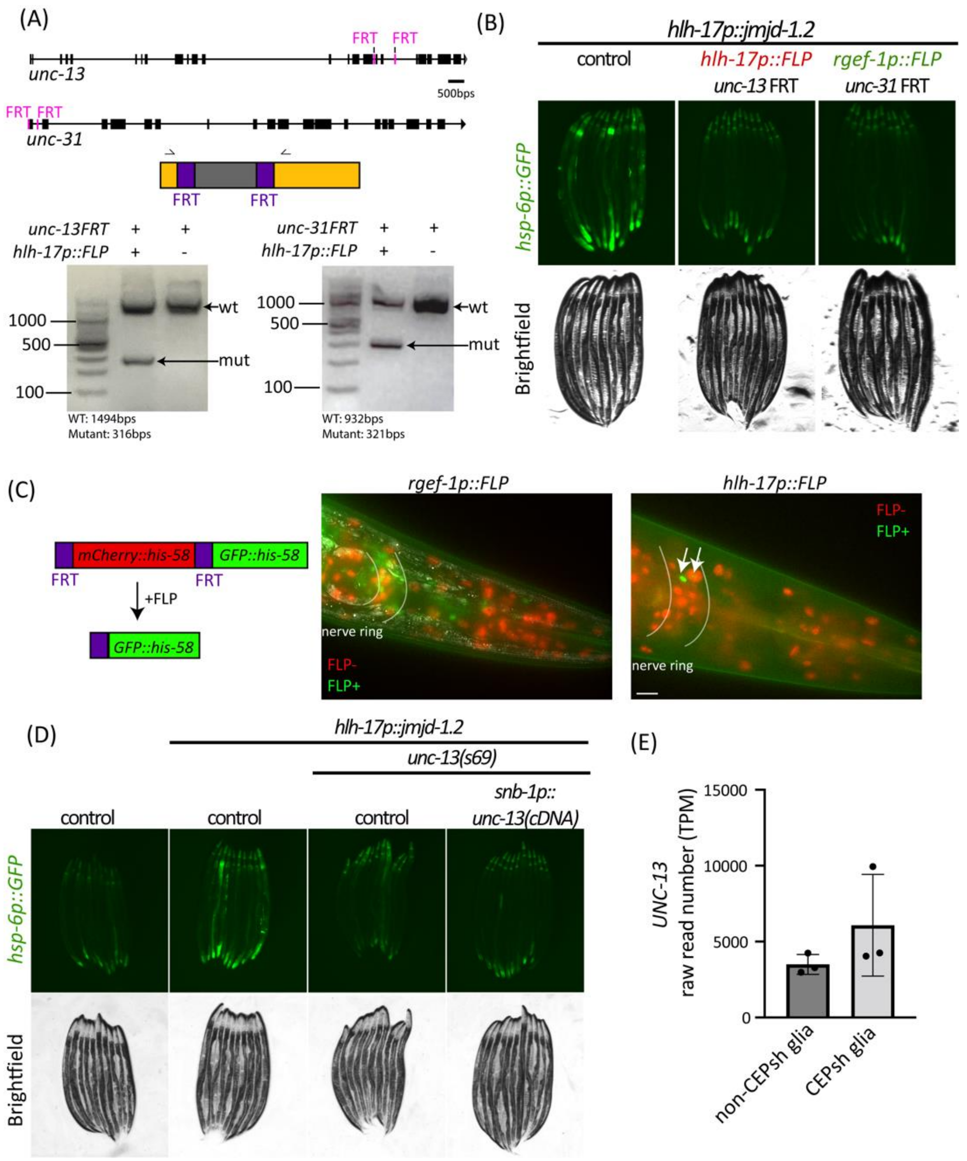


**Figure S5: Cell non-autonomous activation by glial *jmjd-1.2* depends on WNT signaling, related to Figure 5.**

(A) Verification of cutting of newly generated FRT-strains for *unc-13* and *unc-31* (top scheme, FRT location in pink) was verified using PCR of whole-animal DNA extract, resulting in a shorter (cut) fragment when the FLP D5 was present. In addition, all strains were sequenced using Sanger sequencing to verify cutting. (B) Representative fluorescent micrographs as in Figure 5. (C) Verification of the tissue-specificity of FLP alleles (*rgef-1p, hlh-17p*) used in this study was done using a reporter strain, which switches between red-nuclei (FLP negative) and green nuclei (FLP positive) (Muñoz-Jiménez et al., 2017). Arrows indicate CEPsh glia. (D) Rescue of *unc-13* in all neuronal cells could not rescue the activation of the UPR$^{MT}$ in the intestine. (E) UNC-13 read number from CEPsh glia isolated from *C. elegans* (Katz et al., 2019).

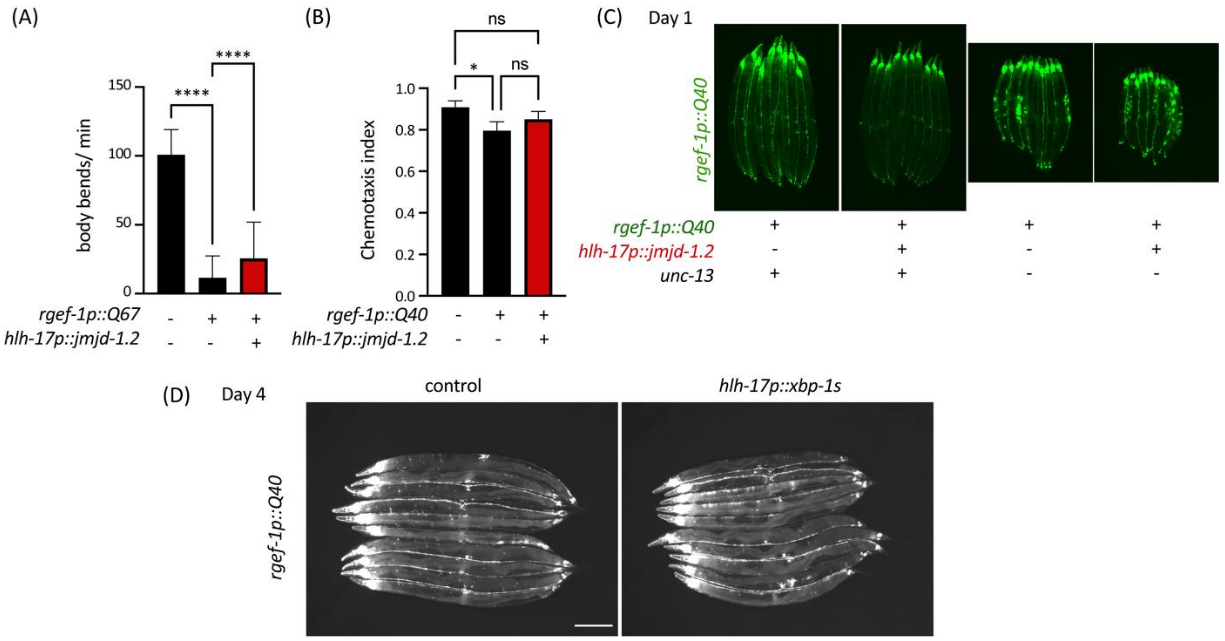


**Figure S6: Glial *jmjd-1.2*, but not glial *xbp-1s*, rescues protein aggregation in neurons, related to Figure 6.**

(A) Thrashing of worms harboring the longer polyglutamine tract, Q67, in neuronal cells, as measured using the WormLab worm tracker (n>50). (B) Chemotaxis as in Figure 7B, towards diacetyl. (C) Same as in Figure 7D, for worms on D1 of adulthood. (D) As in Figure 7D, for worms expressing the ER-UPR transcription factor *xbp-1s* under the *hlh-17* promoter (Frakes et al., 2020). One-way analysis of variance (ANOVA) Tukey's multiple comparisons test, n.s. non-significant, *P < 0.05, ****P < 0.0001.
